# Auditory memory of complex sounds in sparsely distributed, highly correlated neurons in the auditory cortex

**DOI:** 10.1101/2023.02.02.526903

**Authors:** HiJee Kang, Patrick O. Kanold

**Author notes:** **Corresponding author:** Patrick Kanold, Ph.D., Dept. of Biomedical Engineering, Johns Hopkins University, 379 Miller Research Bldg., Baltimore, MD 21205 USA.

## Abstract

Listening in complex sound environments requires rapid segregation of different sound sources e.g., speakers from each other, speakers from other sounds, or different instruments in an orchestra, and also adjust auditory processing on the prevailing sound conditions. Thus, fast encoding of inputs and identifying and adapting to reoccurring sounds are necessary for efficient and agile sound perception. This adaptation process represents an early phase of developing implicit learning of sound statistics and thus represents a form of auditory memory. The auditory cortex (ACtx) is known to play a key role in this encoding process but the underlying circuits and if hierarchical processing exists are not known. To identify ACtx regions and cells involved in this process, we simultaneously imaged population of neurons in different ACtx subfields using in vivo 2-photon imaging in awake mice. We used an experimental stimulus paradigm adapted from human studies that triggers rapid and robust implicit learning to passively present complex sounds and imaged A1 Layer 4 (L4), A1 L2/3, and A2 L2/3. In this paradigm, a frozen spectro-temporally complex ‘*Target*’ sound would be randomly re-occurring within a stream of random other complex sounds. We find distinct groups of cells that are specifically responsive to complex acoustic sequences across all subregions indicating that even the initial thalamocortical input layers (A1 L4) respond to complex sounds. Cells in all imaged regions showed decreased response amplitude for reoccurring Target sounds indicating that a memory signature is present even in the thalamocortical input layers. On the population level we find increased synchronized activity across cells to the Target sound and that this synchronized activity was more consistent across cells regardless of the duration of frozen token within Target sounds in A2, compared to A1. These findings suggest that ACtx and its input layers play a role in auditory memory for complex sounds and suggest a hierarchical structure of processes for auditory memory.

## Introduction

Auditory perception in complex environments requires rapid segregation of different speakers from each other, speakers from other sounds, or different instruments in an orchestra and adjust auditory processing depending on varying sound conditions. Thus, fast encoding of incoming inputs and identifying and adapting to reoccurring sounds is necessary for high-efficiency auditory encoding. This adaptation process can be thought of as an early phase to develop implicit learning of the spectro-temporal stimulus statistics and represents a form of auditory memory (Agus et al., 2010; Luo et al., 2013; Kang et al., 2017). The auditory cortex (ACtx) is known to play a key role on this encoding process. Decreased activity of neuronal responses for repeatedly presented sound is referred as an index of sensory learning (Chew et al., 1995; Chew et al., 1996; Sharpee et al., 2011; Lu et al., 2018). For example, experimental paradigms such as repeated presentation of simple tones have revealed a role of the ACtx in stimulus specific adaption (SSA) (Ulanovsky et al., 2004; Nelken, 2014; Yarden and Nelken, 2017; Yaron et al., 2020). However, such simplified experimental paradigms that are commonly used previously to trigger adaptation of neuronal responses (Ulanovsky et al., 2004; Naatanen et al., 2005; Nelken and Ulanovsky, 2007; von der Behrens et al., 2009) are not sufficient to fully explain processing of spectrotemporally complex acoustic streams that are presented intermittently in dynamic and rich sensory environments. Moreover, the cortical circuits contributing to implicit learning are unknown. In particular, how changes in neuronal responses are established at the population level during the encoding process has not been studied. Higher-order processing would likely require a combination of neural activity across subregions of the ACtx (Ulanovsky et al., 2004; Winkowski and Kanold, 2013; Kanold et al., 2014; Norman-Haignere and McDermott, 2018). However, it is unclear if each ACtx subregion shows distinctive characteristics during the adaptation process and whether the encoding processing follows a hierarchical order across laminae within an ACtx subregion and across ACtx subregions, especially for complex sounds (Wang et al., 2020; Zeng et al., 2021).

To answer the question of how complex dynamic sounds are processed, we acquired neuronal responses simultaneously from many neurons in different ACtx subfields using in vivo 2-photon imaging in awake mice. We adapted an experimental stimulus paradigm from human studies (e.g., (Agus et al., 2010; Andrillon et al., 2015) that triggers rapid and robust implicit learning to passively present complex sounds. In this paradigm, a frozen *‘Target’* complex sound would be randomly reoccurring, instead of consecutive representation of more simplified format of sound exemplars (e.g., tones) that are more commonly used (e.g., (Ulanovsky et al., 2004; Naatanen et al., 2005; Nelken and Ulanovsky, 2007; von der Behrens et al., 2009). While human subjects were only informed to detect whether newly generated abstract random sounds (e.g., broad-band white noise snippets) contained within-sequence repetition for each trial, one specific frozen Target sound re-occurred across trials. Thus, the task does not necessarily drive specific attention only to Target sounds that may lead explicit memory processes, nor are subjects necessarily able to recognize the existence of re-occurring Target sound. Subjects’ task performance level for newly generated sounds versus reoccurring Target sound was used as an indirect measure of implicit learning for Target sound. A series of studies observed a rapid increase of performance specific to presentations of Target sounds, within its 5 intermittent presentations (Agus et al., 2010; Luo et al., 2013; Andrillon et al., 2015; Kang et al., 2017).

By measuring hundreds of neurons simultaneously using in-vivo 2-photon imaging, from three subregions, A1 Layer 4 (L4), A1 L2/3, and A2 L2/3, we identified relevant neural mechanisms of each subregion. We observed a distinct group of cells that are specifically responsive to complex acoustic sequences across all subregions. A decreased amplitude of neuronal responses for Target sounds and more synchronized activity across cells was observed as a memory signature even from a deep layer of A1 where direct thalamocortical input is projected. We also observed more consistent synchronized activity across cells regardless of a duration of frozen token within Target sounds in A2, compared to A1. These findings suggest that ACtx plays a crucial role in auditory memory for complex sounds and subregions within ACtx show both similar and distinctive characteristics on changes in neuronal responses for auditory memory.

## Methods

All procedures were approved by the Johns Hopkins University Animal Care and Use Committee.

### Subjects

We used the F1 offspring of a cross between CBA (Jax#000654) x Thy1-GCaMP6s (C57BL/6J background, Jax#024275). These mice express GCaMP6s in excitatory cells across layers of A1 (Liu et al., 2019; Bowen et al., 2020; Liu and Kanold, 2021) and retain good peripheral hearing (Frisina et al., 2011; Bowen et al., 2020). In humans, clear evidence on learning abstract and complex acoustic sequences is present in young adults (e.g., (Agus et al., 2010; Luo et al., 2013; Andrillon et al., 2015; Kang et al., 2017). Thus, we used young adult mice (N = 17; 8 males, 9 females) between 2-4 months old. Mice were housed in a 12h reverse light/dark cycle room and imaging was performed during the dark cycle.

### Surgery

Surgery was performed as previously described (Francis and Kanold, 2017; Francis et al., 2022). About 1 hour prior to surgery, we injected dexamethasone (1mg/kg, VetOne) subcutaneously (s.c.) to minimize brain swelling. Anesthesia was induced with 4% isoflurane (Fluriso, VetOne) with a calibrated vaporizer (Matrix VIP 3000) and then maintained at 1.5% - 2%. Body temperature was monitored throughout the surgery to maintain around 36C. Hair on the head was first shaved with a shaver and small scissors, and further removed with hair removal product (Nair). Betadine and 70% ethanol were applied 3 times on the exposed skin. Then skin and tissues were then removed and muscles were scraped to the left temporal side. Unilateral craniotomy was performed to expose about 3.5 mm diameter region over the left ACtx. Three circular glass coverslip (2 of 3 mm and one 4mm coverslips) was affixed with a clear silicone elastomer (Kwik-sil, World Precision Instruments). A custom 3D-printed stainless steel headpost was attached on the skull and the rest of exposed skull area and the edge of the cranial window were covered using a dental acrylic (C&B Metabond). Carprofen (5mg/kg), cefazoline (300mg/kg), and additional dosage of dexamethasone (1mg/kg) were injected (s.c.) post-operatively. Animals had 7-10 recovery days before any imaging was performed.

### Stimuli and paradigm

All acoustic stimuli were pre-generated using Matlab (Mathworks, version R2020b), calibrated at 70dB SPL using a Bruel & Kjaer 4944-A microphone. Stimuli were loaded using Tucker-Davis Technologies (TDT) RX6 processor for loading sounds and PA5 attenuator to adjust sound levels, and delivered via ES1 speaker using ED1 speaker driver.

Pure tones: A series of pure tones in different frequencies and sound levels were played to 1) define subregions within ACtx using widefield imaging before 2-photon Ca^2+^ imaging, and 2) distinguish each cell’s tuning property during 2p imaging session, in addition to the experimental imaging session playing complex sounds. For widefield imaging, 100 ms pure tones in different frequencies from 4 kHz to 64 kHz were presented in three sound levels (30, 50, 70 dB SPL). For 2p imaging, 100 ms pure tones from 4 kHz to 54 kHz (16 log-spaced frequency grid) were presented in two different sound levels (40, 60 dB SPL) to keep each imaging session not to go over 1 hour to minimize a risk of photobleaching and discomfort of animals. From these stimuli we determined the frequency response are (FRA) of cells (2P imaging) or ROIs (WF imaging).

Dynamic Random Chords (DRCs): Adapted from previous studies (Agus et al., 2010; Luo et al., 2013; Andrillon et al., 2015; Kang et al., 2021), we presented a series of spectro-temporally complex artificial sounds, dynamic random chords; DRCs. One segment of DRC is defined as about 200-ms duration in which at each time bin (20 ms) filled with equivalent length of a tone pip with varying sound levels (50 – 90 dB SPL) at each frequency bin (4-40 kHz separated by 20 frequency bins following a logarithmic scale) (**Fig. 1A, left**). Each tone pip within the time bin had 5 ms of rise and fall; fall-ramping of the preceding tone was overlayed with rising ramping of the following tone, thus the actual duration of each segment became about 180 ms. One sequence of DRC consisted of 5 segments, in a format of two different types. First, a 180 ms segment was concatenated for 5 times, containing within-sequence repetition (Random_rep_). The second type was generated by 5 different segments concatenated as a fully-random long complex sound (Random_long_). 50 different sequences for each type were generated, referred as Random conditions. In addition, one additional sequence in each type was generated as frozen exemplar to be presented 50 times intermittently, referred as Target conditions (Targe_trep_ and Target_long_). By having these four combinations across types (rep. or long) and conditions (Random or Target), we aimed at identifying how the duration of frozen segment within Target sound affects the encoding speed, across subregions. Lastly, all sequences had additional pre- and post-random DRC segment (180 ms). This was to minimize any sound onset or offset-driven effect to interfere with response changes during the actual sequence presentation. In total, each test block had 200 trials, each trial presenting each DRC at a random order (**Fig. 1A, right)**. The random order of sequence presentation was pre-allocated and applied in the same order across animals so that there is no variability of sequences at a single trial level across subjects. A total of 15 test blocks was pre-generated and randomly selected test block was played for an imaging session. While multiple imaging sessions across days were performed per animal, no animal received the same test block. Two blocks containing different stimuli of DRCs were performed in one imaging session. Between two DRC blocks, a FRA block was presented to acquire tuning property of cells.

**Figure 1.**
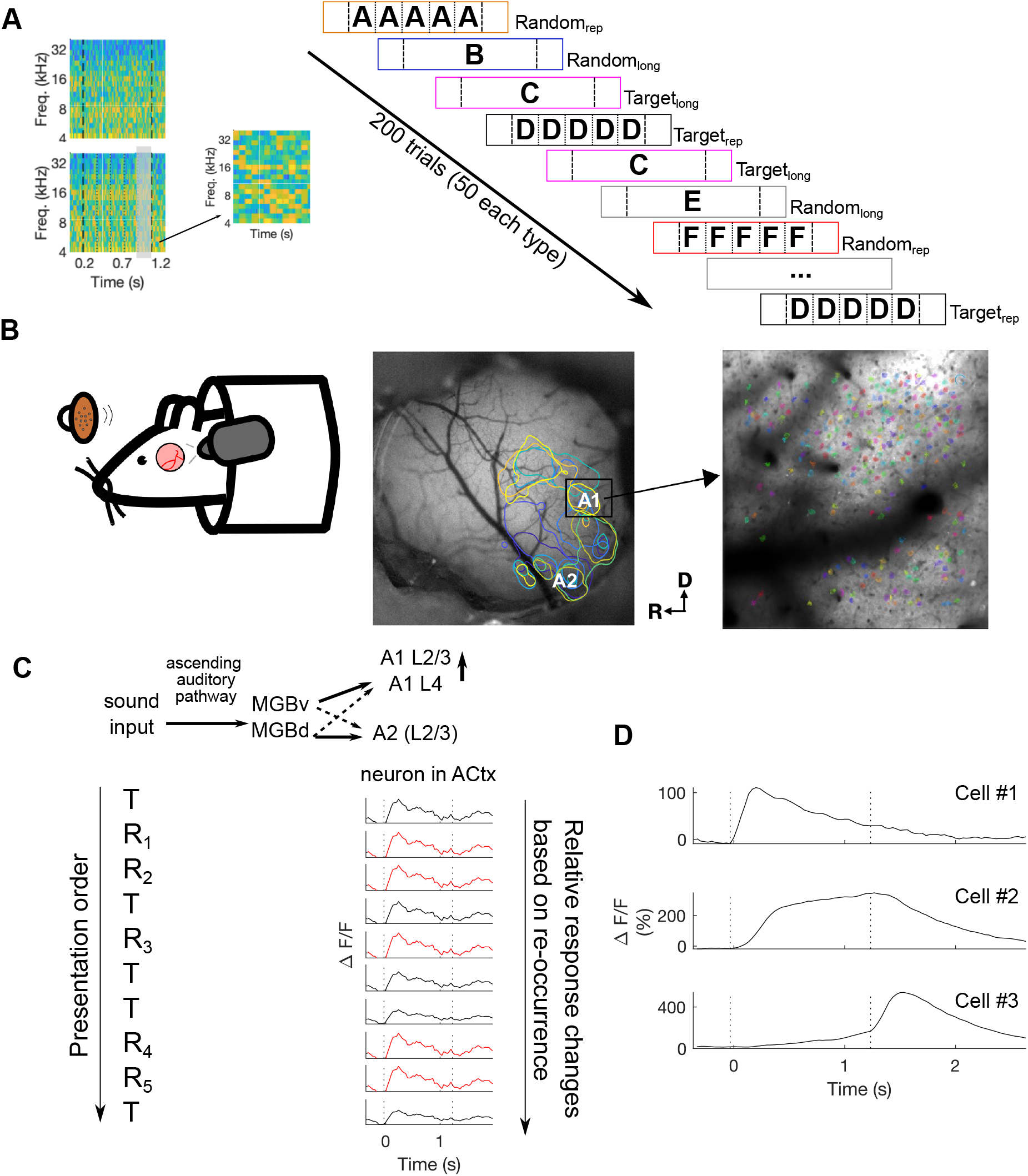
Experimental approach using two-photon imaging in the ACtx using Thy1-GCaMP6 transgenic mice. **A**: (left) Example spectrogram of DRC. One sequence of DRC is consisted of 20 ms tone pips in each frequency and time grids at different sound levels, for about 1.2 sec. DRC sequence contain either different segments (top; long sequence) or the same segment multiple times (bottom; rep. sequence). Black dashed vertical lines indicate onset and offset of DRC sequences where stimulus condition will be defined. Black dotted vertical lines indicate 180 ms length segment embedded 5 times generating within-sequence repetition for rep. sequences. The inset on the right is a magnified spectrogram of the segment marked in gray. The sequence is further calibrated at 70 dB SPL for each frequency grid and overall sound level. (right) Example schematic of stimuli presentation during a recording session. One imaging session includes a series of either newly generated DRCs (Random) or one specific re-occurring sequence (Target). All stimuli have pre- and post-random segments. Sequences with the same color codes *<u>and</u>* the same letter refer to Target sequences (Target_rep_ and Target_long_) while sequences different color codes refer to Random sequences (Random_rep_ and Random_long_) generated afresh for each presentation. Long sequences are about 0.9 sec. DRC, while rep. sequences are 0.18 sec., with additional 0.18 sec. of pre- and post-random DRC segments regardless of conditions. **B**: We image over the cranial window of the ACtx while a head-fixed mouse receives a passive exposure to DRCs. After locating the ACtx based on a tonotopic map acquired by wide-field imaging as left example image (low to high frequency marked with darker to lighter color. R: rostral, D: dorsal), two-photon imaging is conducted to trace cell responses in more focused areas such as A1 or A2. Right example image indicates hundreds of imaged cells in A1. **C**: An example schematic of traces of neuronal responses for passive sequence presentation. Decreasing amplitude of sound evoked response to Target sound (T; black) from its re-occurrences, as early as its second presentation, compared to constant sound evoked neuronal responses to Random sequences (R1, R2, … R5; red) is regarded as an index of ‘learning’. **D**: Example fluorescence trace of selected neurons as DRC responsive from A1 L2/3 imaging. Black dashed lines indicate sound onset and offset.

### Widefield imaging

Widefield imaging was performed as previously described (Liu et al., 2019). Animals were restrained in a 3D printed plastic tube and head fixed on a custom designed headpost holder. In-vivo widefield imaging was performed by sending 470nm LED light (Thorlabs, #M470L3) over ACtx using a camera (PCO edge 4.2) to capture 330 x 330 pixel sized images covering the 3mm diameter cranial window at 30 Hz frame rate. A series of 100 ms pure tones between 4 - 64 kHz in 30, 50, 70 dB SPL were presented at a random order to identify regions of ACtx.

### 2-photon imaging

In-vivo 2p Ca2+ imaging was performed over ACtx while animals are head fixed using Bruker 2P Plus microscope tilted at 48~55 degrees to target the ACtx with 940 nm excitation wavelength (Coherent Discovery), Nikon LWD 16x Objective (0.80 NA, 3.0 WD), and PrairieView software (version 5.6). The field of view size was about 512 x 512 μm and imaging frame rate was 30 Hz. For A1 L2/3 and A2 L2/3, we imaged at a depth between 160-180 um, and for A1 L4, we imaged at a depth of ~410 um (Winkowski and Kanold, 2013).

#### Data analyses

##### Widefield imaging

Image processing followed a method reported in the study of (Liu et al., 2019). Briefly, on a downsampled image sets by a factor of 3, we applied an autoencoder neural network to run an automatic image segmentation to group pixels with strong temporal correlations as a single component (Regions of Interests, ROIs) by applying a dimensionality reduction (Whiteway and Butts, 2017) and pulled out 50 ROIs. For each ROI, images of 15 frames from the sound onset (F_t_) were baseline corrected with average value of 5 frames before each onset to trace fluorescence changes to different frequency-sound level pairs (F_norm_ = (F_t_ -F_0_)/F_0_). We defined subregions of ACtx from a tonotopic gradient of tone responses for each subregion (Liu et al., 2019). Here we focus on A1 and A2. For A1, a low-to-high frequency gradient (tonotopy) was organized following the caudal to dorsomedial direction. For A2, the tonotopy was reversed, following the medial to ventrolateral direction (**Fig. 1B**).

##### 2-photon imaging

Motion correction, cell detection and cell fluorescence traces including neuropil fluorescence were extracted from raw imaging data using suite2p software (Pachitariu et al., 2016). Among detected cells, we selected cells that show sound-evoked responses, following our previously used method (Liu et al., 2019; Liu and Kanold, 2021). Neuropil correction was computed first following the equation of: F_(cell_corrected)_ = F_(cell)_ – (0.8 * F_(neuropil)_). Then we computed ΔF/F by first subtracting the average baseline period before the sound onset (100 ms) from F, dividing by the baseline, then multiplying by 100 to denote the change values in %. To select cells that are sound responsive, we compared 99% confidence interval of baseline (100 ms before the sound onsest) and sound-evoked activity (1 sec after the sound onset). Cells were regarded as sound responsive when the lower bound of CI during the sound presentation is above the upper bound of CI during baseline (**Fig. 1D**). About 20 – 50% of cells per imaging session were selected as sound-responsive cells and used for further analyses. We then divided DRC responsive cells into subgroups based on the sound evoked response amplitude difference between Target and Random conditions, as we aimed at tracing adaption of cell responses (i.e., decrease in evoked response amplitude) as an index of learning for Target sounds (**Fig. 1C**).

##### FRA cell classification

Based on tonal responses from 2-photon imaging, we classified cell shapes, following a previously used method from (Liu and Kanold, 2021). Briefly, we aligned the FRA of all tonal responsive cells at the geometric center of the FRA, with the weighted average of frequencies by significant responses, followed by a principal component analysis (PCA) to reduce the dimensionality to explain 95% of the variance in the aligned FRA. K-means clustering (number of clusteres = 5) and T-distributed stochastic neighbor embedding (t-SNE) algorithm were then applied to obtain the clustering result. Five clusters include classic H, V, I-shaped FRAs and two sparsely responding shapes (S1 and S2; **Fig. 2A**).

**Figure 2.**
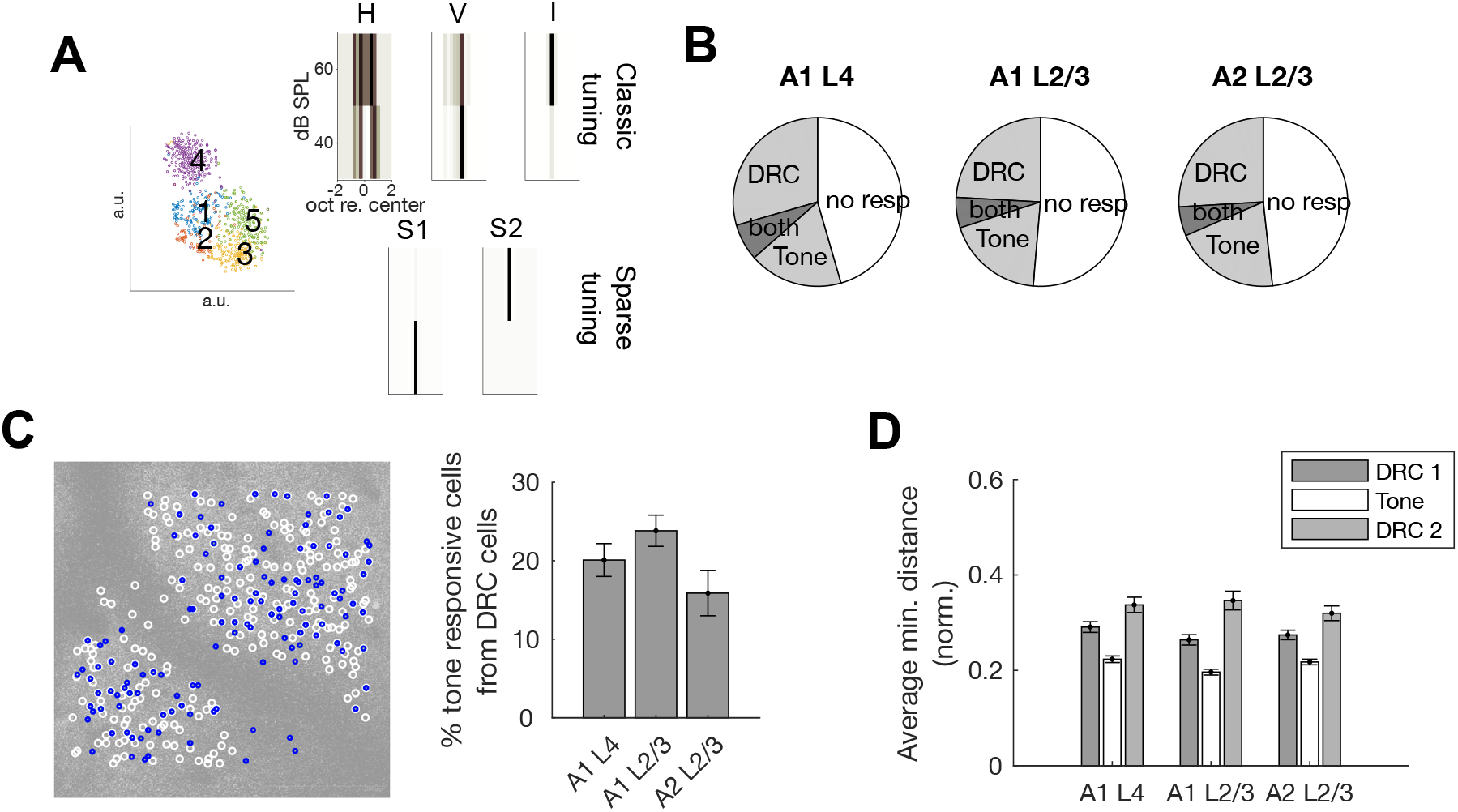
**A**: Example distribution of cells responsitve both for DRC and FRA in A1 L4 clustered into 5 different categories. **B**: Cell distribution per stimulus type. Proportion of sound-responsive cells (about 49 – 55% of cells; grey) for each subregion. Only about 6-7% of cells were responsive for both types of sounds. **C**: An example distribution of either DRC (white) or FRA (blue) responsive cells on an imaged field of view (512 x 512 um) in A1 L2/3. About 20% of DRC cells also respond to pure tones. Errorbars indicate standard error of mean (SEM) across imaging sessions. **D**: Normalized average minimum distance between responsive cells for DRC (gray bars) and pure tones (white bar). Errorbars indicate SEM across cells.

##### Signal/noise correlation

Signal correlation are the synchronised activity between neurons for each stimulus type. We computed cross-correlation on average ΔF/F traces across trial, for each stimulus type, between two neurons at zero lag using matlab function ‘xcorr’ (Winkowski and Kanold, 2013). To measure similarity on trial-to-trial fluctuation between neurons per stimulus type as noise correlation, we applied the same cross-correlation function on a vector of mean amplitudes during the sound presentation per trial level between two neurons for each stimulus type, using matlab function ‘xcov’. We compared the distribution of signal/noise correlation indices between Target and Random sounds for both short and long sequences.

##### Statistical analyses

For comparisons on the proportion of cells that are responsive to both DRC and tones, we ran one-way analysis of Variances (ANOVA). For multiple comparisons of the minimum distance between sound types (DRC or FRA), we ran one-way ANOVA across subregions and post-hoc one-way ANOVA for each subregion across sound types. For comparisons of fluorescence traces across stimulus types (Short and long Target or Random DRCs), cluster-based permutation paired *t*-tests on short Target versus Random and long Target versus Random across cells as independent observations were performed, with 1000 iterations (*alpha* < 0.01). To compare neuronal synchrony based on stimulus type, we computed repeated measures (RM ANOVA) as each neuron pair as random factor, two factors in conditions (duration of segment either repeating or long, and re-occurring target or random) as independent variables, and SCs as dependent variables. Paired *t*-tests on signal and noise correlation between Target and Random for short and long cases respectively were computed post-hoc. Significance level was Bonferroni-corrected for multiple comparisons.

## Results

We sought to identify if cells in ACX showed a neural signature of implicit memory and where in the cortical hierarchy this signature emerged. We first defined the subregions within ACtx by acquiring frequency responsive area (FRA) maps from widefield imaging while a series of pure tones in different frequencies (4-64 kHz) were played (Liu et al., 2019; Romero et al., 2020) (**Fig. 1B**). We then presented a series of DRCs (**Fig. 1A**) in passively listening awake mice while conducting two-photon imaging in three different subregions: thalamocortical layer L4 of primary auditory cortex (A1), superficial layer L2/3 of A1, and superficial layer L2/3 of the secondary auditory cortex (A2). A series of different DRCs, referred as ‘*Random*’ sounds, were generated in two ways. They were either a 180 ms random DRC segment concatenated for 5 times including within-sound repetition (Random_rep_), or equivalent length (i.e., 900 ms) of DRC segment (Random_long_). Among a series of random sounds of either type, one specific sound, referred as *‘Target’* sound, was generated in either way (Target_rep_ or Target_long_) kept the exact spectrotemporal characteristics and re-appeared at random trials. Thus, one imaging session contained 4 conditions (Random_rep_, Random_long_, Target_rep_, and Target_long_) in total, with each condition of 50 trials, presented in a randomized order. We designed the stimulus paradigm to minimize the effect of simple adaptation to repeating simple auditory stimuli. Thus, any changes observed in sound-related fluorescence traces between Target and Random sequences was closely correlated to the encoding of re-occurring (Target) input.

### Cells in A1 and A2 respond to tones and DRCs

We sought to investigate changes in response amplitudes elicited by the Target sound from its random re-occurrences, compared to Random sounds, regardless of the duration of the frozen token. We measured fluorescence changes of 8085 ACtx cells (2929 cells in A1 L4, 3510 cells in A1 L2/3, and 1646 cells in A2 L2/3) by two-photon imaging while mice were passively listening to pure tones, and the series of acoustically complex and abstract auditory patterns (DRCs) of 4 different conditions presented at a random order as explained above.

We first investigated if subregions of ACtx showed a preference to tonal stimuli or to DRCs. We computed the average sound-evoked activity during the presentation of pure tones generating a frequency response area (FRA) for each cell and also computed the responsiveness during DRCs. We found that between 15-19% of cells respond to pure tones, while 31-45% respond to DRCs irrespective of subregions (**Fig. 2B**). We found that about 16-20% of DRC cells also respond to pure tones across all subregions (*F*(2,62) = 2.64, *p* = 0.08; **Fig. 2C**). DRCs consisted of a wide frequency range covering from 4 kHz to 40 kHz, and we thus investigated if cells with certain best frequency (BF) were more responsive to DRCs. Cells of all BFs were responsive for both DRCs and tones across all subregions with an average BF of around 15 kHz. To identify if tone and DRC responsive cells showed a different spatial distribution, we next measured the minimum distance between each cell pair for each sound type, for each subregion. We observed that in all imaged subregions the distribution of cells responsive to either DRC or tones is rather independent, with a greater minimum distance between cells observed for DRC compared to pure tones, suggesting a sparser distribution of DRC cells (*F*(2,5518) = 3.11, *p* = 0.045 across subregions; *F*(2,1976) = 31.47, *p* < 0.0001 across sounds for A1 L4, *F*(2,1447) = 47.07, *p* < 0.0001 across sounds for A1 L2/3; *F*(2,2089) = 30.34, *p* < 0.0001 across sounds for A2 L2/3; **Fig. 2D**). Data shows that cells in both layers of A1 and A2 are responsive to both tones and DRCs. Moreover, cells responsive to more complex sounds are more sparsely distributed regardless of subregions. For any further analyses, we only focused on cells that were DRC responsive.

### Sound-evoked activity decreases for re-occurring sounds regardless of sequence duration

To identify if a cell’s responsiveness depends on the sound reoccurrence, we examined the differences of the sound-evoked fluorescence changes from the sound onset (ΔF/F) between Target and Random sounds. Our stimuli contained two Target conditions: Target_rep_ and Target_long_. In the ‘repeat’ version the 180 ms Target segment is concatenated for 5 times while the long version comprises a single 900 ms Target segment. Thus, a difference in the response to these two conditions would indicate that cells are able to respond differentially based on the duration of frozen token (180 ms versus full 900 ms) and within-sequence repetition.

We observed a generally larger response of the Random sound over the re-occurring Target sound in all three subregions of ACtx, especially for the Target_rep_ sound (**Fig. 3A**). In particular, we observed that a fraction of cells was showing decreased sound-evoked activity for both short and long Target sounds in all subregions (29% for A1 L4, 24% for A1 L2/3, and A2 L2/3; **Fig. 3B)**. More sustained activity to Target_rep_ compared to Target_long_ indicates that cells are sensitive to within-sequence repetition. These data indicate that subsets of ACtx cells are sensitive to the re-occurrence of complex sounds and that such cells are already present in the thalamocortical input layer L4 of A1. Since targets and Random sounds overlap in the spectral domain, this indicates that ACtx cells can rapidly learn the identity of complex sounds.

**Figure 3.**
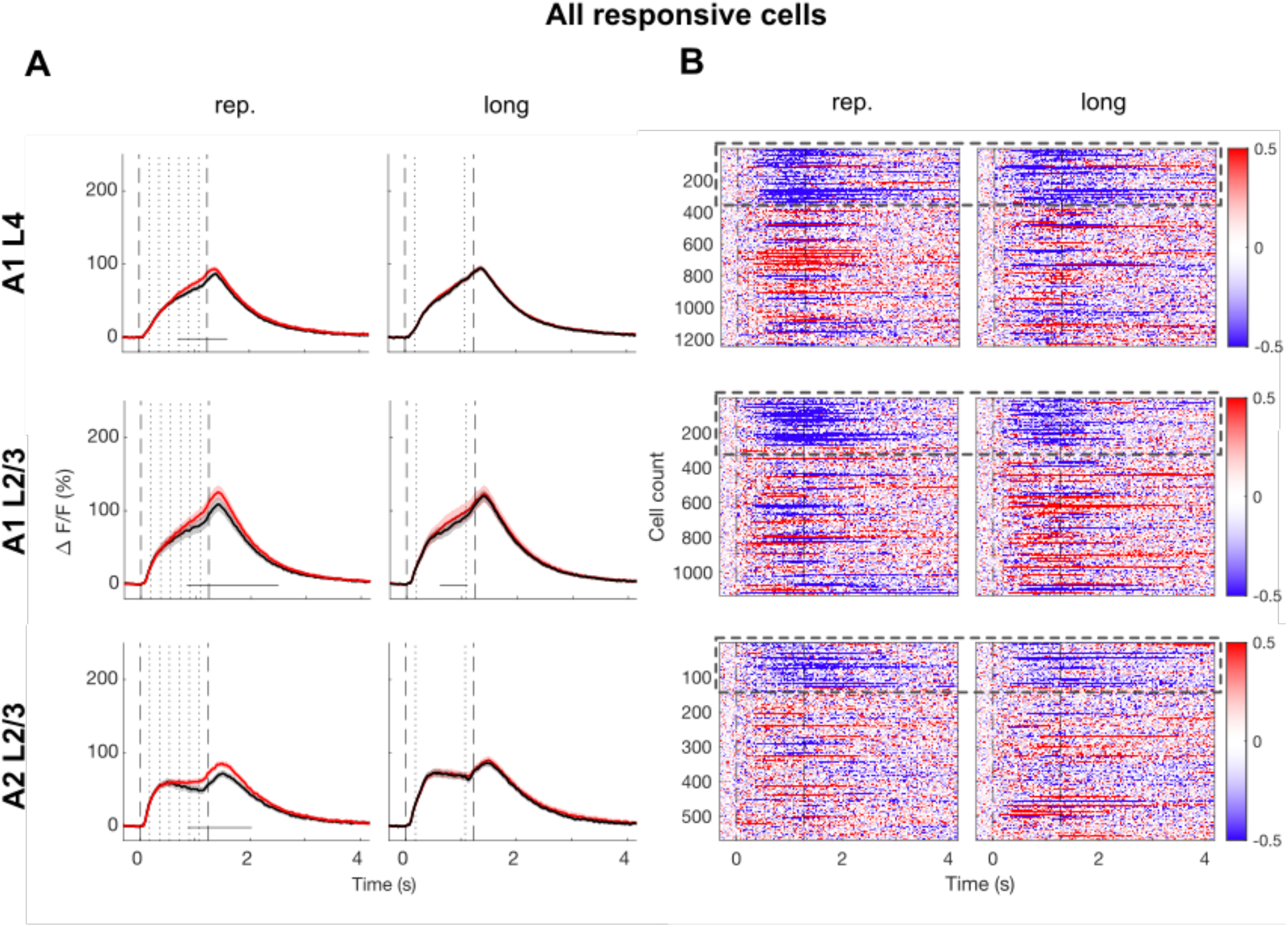
**A**: Traces show the average evoked fluorescence change (Δf/f) in neuronal populations in each cortical region (top: A1 L4, middle: A1 L2/3, bottom A2 L2/3) to the sound presentation (black: Target, red: Random; left column: rep. sequences, right column: long sequences) from two-photon imaging. Dashed vertical lines indicate the sound onset and offset of the entire sound including pre and post random segments and dotted vertical lines indicate the target sequence onset and offset. Shaded areas indicate 95% confidence intervals. Horizontal solid lines just above x-axes indicate time points where significant differences between conditions were observed (cluster permutation with 1000 iterations, alpha < 0.01). **B**: Heatmap on a trace difference between Target and Random sequences of each cell activity across subregions. About 24-29% of cells marked with dashed rectangular boxes showed decreased activity for Target sequence regardless of frozen token duration.

To further verify whether such decreased activity is due to re-occurrences of the Target sound, we examined the mean amplitudes of cell activity during the sequence presentation per each trial. We chose to average traces from 5 frames after the sequence onset to 10 frames after the sequence offset to compensate for the slow dynamics of GCaMP6s fluorescence. The average amplitude was more greatly decreased along trials for Target sounds compared to Random sounds, across all subregions (**Fig. 4**). Target_rep_ showed greater difference to Random sounds compared to the Target_long_, possibly due to Target_rep_ containing additional within-sequence repetition.

**Figure 4.**
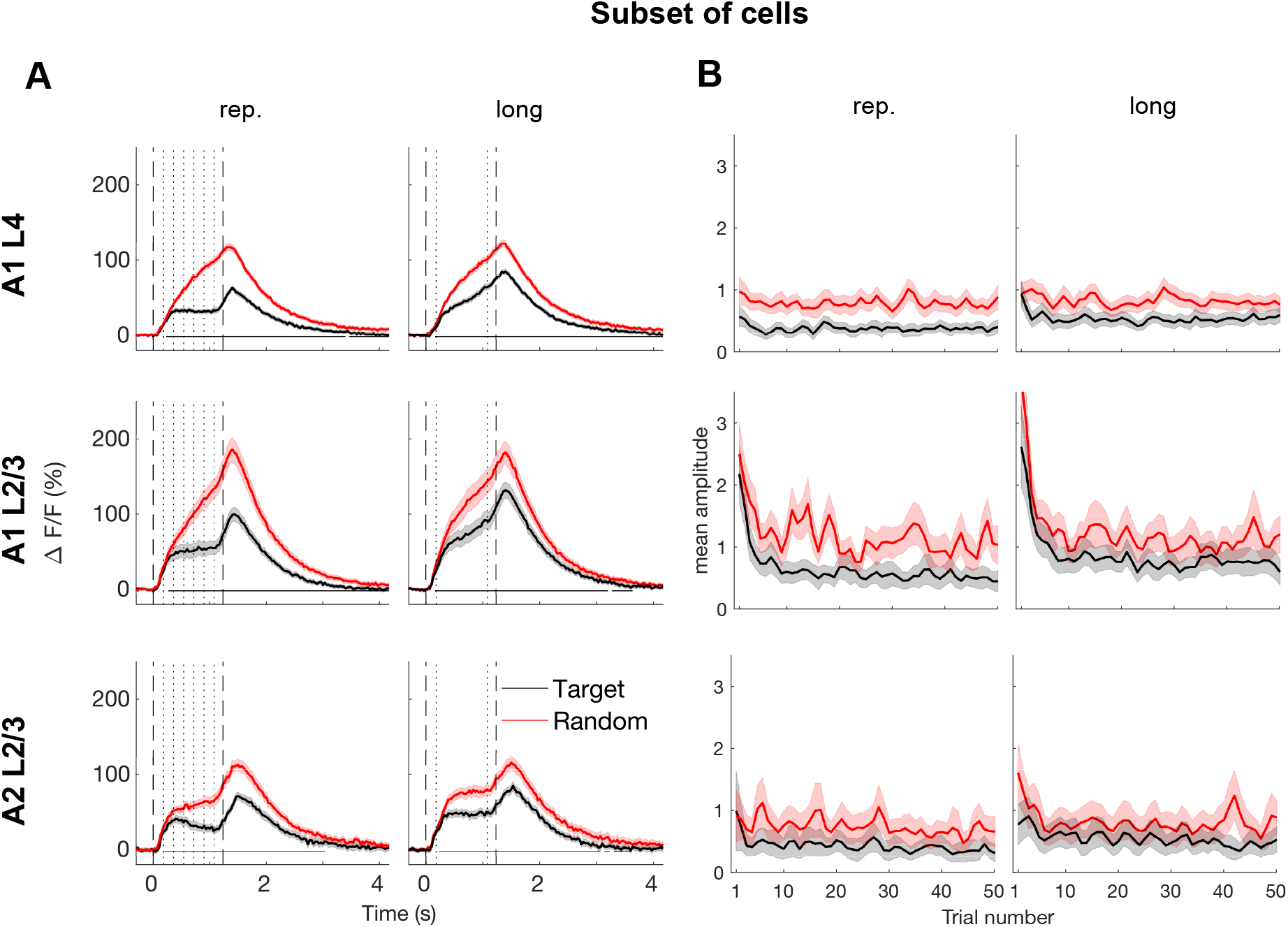
**A**: Average evoked fluorescence change (ΔF/F) in a subset of neuronal populations in each cortical region (top: A1 L4, middle: A1 L2/3, bottom A2 L2/3) to the sound presentation (black: Target, red: Random; left column: rep. sequences, right column: long sequences) from two-photon imaging. Dashed vertical lines indicate the sound onset and offset of the entire sound including pre and post random segments and dotted vertical lines indicate the target sequence onset and offset. Horizontal solid lines just above x-axes indicate time points where significant differences between conditions were observed (cluster permutation with 1000 iterations, alpha < 0.01). Shaded areas indicate 95% confidence interval. **B**: Average amplitude across the subset of cells during sound presentation at a single trial level. Greater decrease on the average amplitude on Target than Random sequences were observed at a rapid pace, as early as trial 2. This indicates adaptation of neuronal responses to re-occurring Target sounds, referred as an index of learning.

### Sensitivity for re-occurring sounds depends on functional cell type

Our sample of DRC responsive cells contained cells that were unresponsive to tones. We thus wondered if there was a difference in the sensitivity of cells to targets between tonally unresponsive and responsive cells. Moreover, since tonally responsive cells show a variety of FRAs indicating different functional cell types (Liu and Kanold, 2021) (**Fig. 2A**), we wondered if Target selectivity differed with functional cell type. We thus applied the same approach to examine activity changes of DRC responsive cells that were also responsive to tones and showed a clear FRA (about 15 - 20% of DRC cells). We divided cells into tuned cells and sparse cells. While few conditions show significant response difference between Target and Random sequences due to limited number of cells that were responsive for both DRCs and FRA, different cell types seemed to show different effects per subregions. First in A1 L4, tuned cells showed significant response difference especially for Target_long_ when compared to Random_long_. In A1 L2/3, on the other hand, it was sparse cells that showed a response difference between Target and Random sequence (Target_rep_ and Random_rep_). In A2, no distinctive difference between cell types was observed (**Fig. 5**).

**Figure 5.**
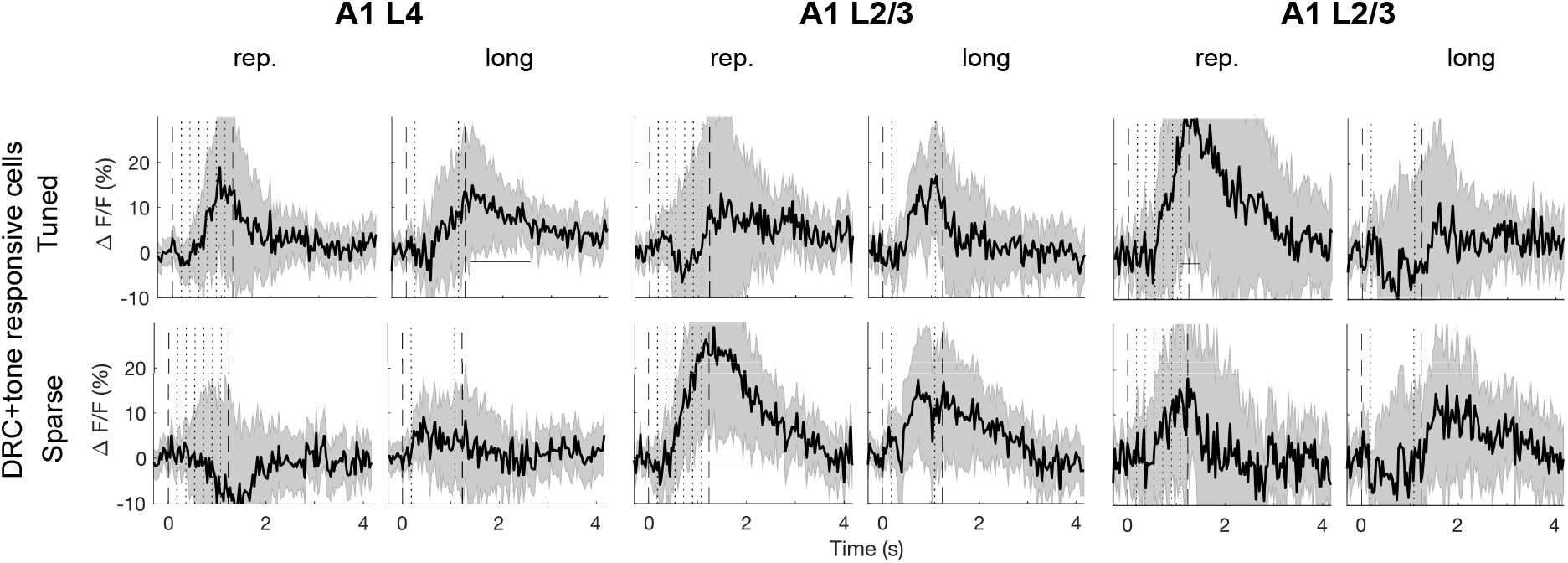
Average response difference between Target and Random sequences (ΔF/F(Random) - ΔF/F(Target)) from a subset of neuronal population where tuning property and cell type could be defined, in each cortical region (left: A1 L4, middle: A1 L2/3, right: A2 L2/3) to the sound presentation (left columns per each region: Random_rep_ – Target_rep_; right columns per each region: Random_long_ – Target_long_). from two-photon imaging. Dashed vertical lines indicate the sound onset and offset of the entire sound including pre and post random segments and dotted vertical lines indicate the target sequence onset and offset. Horizontal solid lines just above x-axes indicate time points where significant differences between conditions were observed (cluster permutation with 1000 iterations, alpha < 0.01). Shaded areas indicate 95% confidence intervals across cells.

### Implicit learning is driven by ensemble activity from a subset of neurons

So far, we have examined changes in the response amplitude of single ACtx cells to DRCs that occur during the learning of new sounds, regardless of sound duration. Since neurons do not encode stimuli in isolation, but as a population (Francis and Kanold, 2017), we next investigated whether activity changes of selective ‘learning neurons’ show greater functional connectivity within the imaging field. Signal and noise correlations are commonly used methods to identify response synchrony and functional connectivity across neurons (Cohen and Maunsell, 2009; Cohen and Kohn, 2011; Winkowski and Kanold, 2013). First, we computed signal correlation (SC) of the responses to the full sound duration of the DRC averaged across trials for each neuron pair per subregion. We then compared SCs between Target and Random conditions. As above, Target and Random conditions were further subdivided into rep. (180 ms segment repeating 5 times) and long (900 ms. segment) to compute and compare SCs to investigate the possible difference in correlations based on the duration of frozen segments.

First of all, differences in SCs across conditions (Target vs. Random) and segment duration (rep. vs. long) were observed in all subregions (Condition - A1 L4: *F*(1, 4834) = 10452.20, p < 0.0001; A1 L2/3: (*F*(1, 4679) = 7224.12, p < 0.0001; A2 L2/3: (*F*(1, 841) = 1277.60, p < 0.0001; segment duration - A1 L4: *F*(1, 4834) = 279.70, p < 0.0001; A1 L2/3: (*F*(1, 4679) = 582.79, p < 0.0001; A2 L2/3: (*F*(1, 841) = 95.74, p < 0.0001). We also observed interactions between condition and segment duration (A1 L4: *F*(1, 4834) = 463.98, p < 0.0001; A1 L2/3: (*F*(1, 4679) = 1146.07, p < 0.0001; A2 L2/3: (*F*(1, 841) = 13.69, p = 0.0002). Post-hoc paired t-tests per each Target - Random pair for each segment duration revealed a difference across subregions. That is, for A1 L4, the SCs to Target_long_ but not Target_rep_ condition were larger than for equivalent Random conditions (*t*(4834) = −0.89, p = 0.37 for Target_rep_ and Random_rep_ pair, *t*(4834) = 20.43, p < 0.0001 for Target_long_ and Random_long_ pair). In A1 L2/3, SCs for Target_rep_ was lower than Random_rep_ while SCs for Target_long_ were greater than Random_long_ (*t*(4679) = −8.14, p <0.0001 for Target_rep_ and Random_rep_ pair, *t*(4679) = 31.51, p < 0.0001 for Target_long_ and Random_long_ pair). Lastly, in A2 L2/3 SCs for both long and short Target were greater than for Random sounds (*t*(841) = 13.12, p < 0.0001 for Target_rep_ and Random_rep_ pair, *t*(841) = 4.01, p < 0.0001 for Target_long_ and Random_long_ pair; **Fig. 6A**). These results indicate that cells across areas show higher SCs for long Targets were larger than for Random sounds.

**Figure 6.**
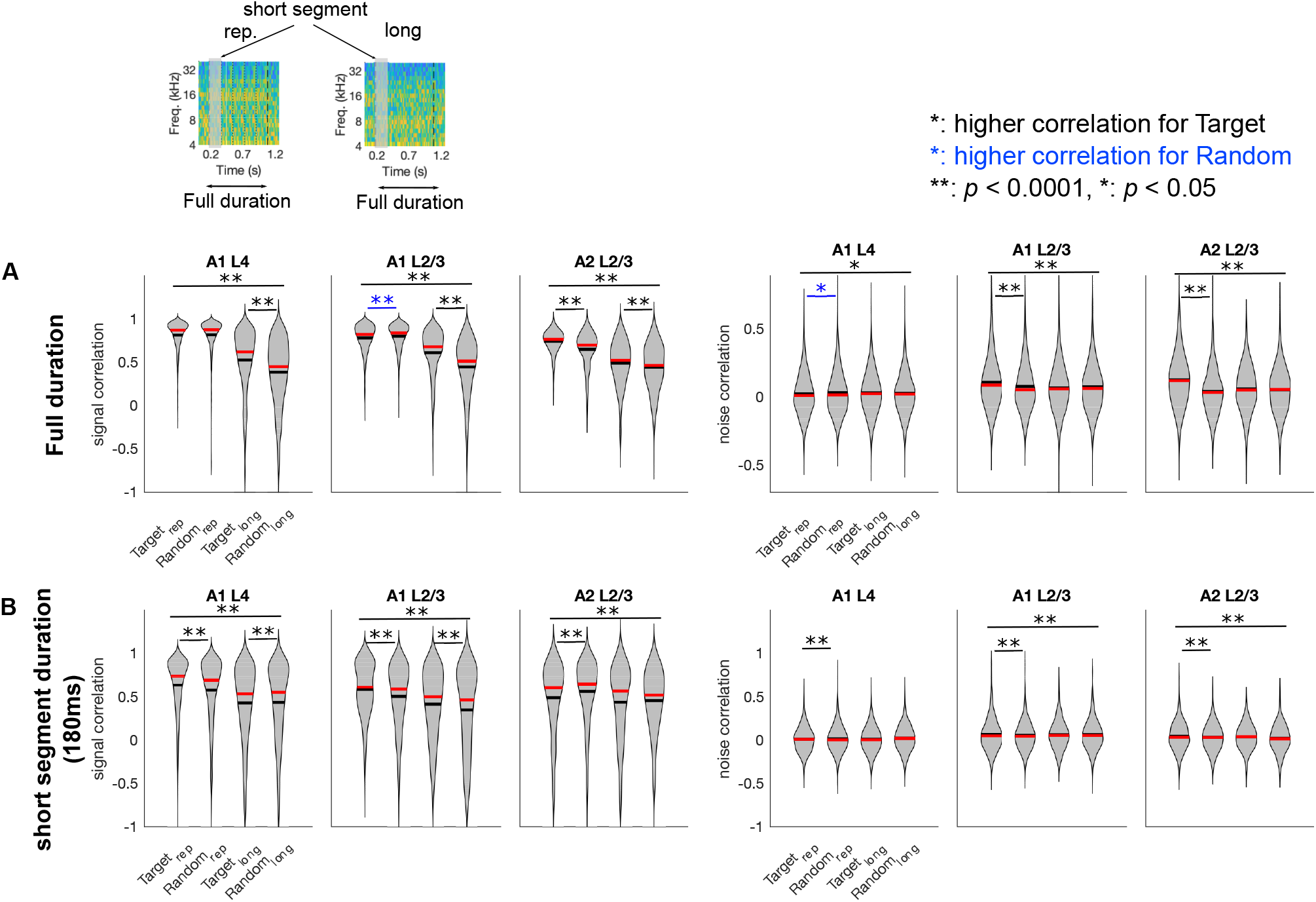
Violin plot of SCs and NCs across cell pairs per each condition and each frozen token duration (rep.: left column, long: right column), for full duration (**A**) and first segment duration (**B**). Top insets are example spectrogram and duration included to compute SCs for each case is marked. Greater correlation for Target_long_ was observed across all subregions only when full duration of sequence was taken into the account. For Target_rep_, significantly greater signal correlation was observed for full duration computation in A2, whereas only first segment duration was required for A1 L4 and L2/3. Black lines on the violin plots are mean values across pairs and red lines are median values. See text for more details.

We speculated that the similarity for short Target versus Random in the deep layer of A1 might be due to A1 being able to process information on shorter (and thus faster) time scales than along the high-order auditory pathway (Lim et al., 2016; Asokan et al., 2021). Both types of stimuli contain within-sequence repetitions for five times. Thus, processing on a rapid timescale in A1 might enable the encoding of the frozen token, even for Random sequences, leading a similarity between two conditions. To further verify this notion, we took a shorter signal trace that is relevant to only the first segment of sequences in the short segment case. If the trace reflects learning over trials specifically for the Target condition, even for the short segment case, there still could be a difference between Target and Random condition in A1. Conversely, for the long case, the distinctive characteristic of Target sounds may not always appear in the beginning of the sequence, thus there may be a less clear difference between Target and Random when the entire trace is not included but limited to only first 180 ms durations, especially for A2 where global feature of frozen segment may be considered for encoding process. As expected, greater SCs were observed especially for short sequences of Target versus Random sounds in A1 for both layers (A1 L4: *t*(5010) = 6.99, p < 0.0001 for Target_rep_ and Random_rep_ pair, *t*(5010) = −2.7961, p = 0.005 for Target_long_ and Random_long_ pair; A1 L2/3: *t*(4817) = 3.01, p = 0.0027 for Target_rep_ and Random_rep_ pair, *t*(4817) = 11.76, p < 0.0001 for Target_long_ and Random_long_ pair; **Fig. 6B**). In A2 L2/3, greater SCs were only observed for Target_rep_ and Random_rep_ pair (*t*(878) = 4.02, p < 0.0001 for Target_rep_ and Random_rep_ pair, *t*(878) = 2.32, p = 0.02 for Target_long_ and Random_long_ pair; *p*-values are uncorrected values and significance threshold was adjusted by Bonferroni correction to compensate multiple comparisons). This shows that neurons in A1 display synchronized activity for both segment duration of Target sounds. Also, the synchrony across neurons in A1 is established in shorter time scale, compared to A2, which yield high SCs even for Random_rep_ cases as they include within-sequence repetitions. Together this suggests that local networks during the encoding process may be shaped differently between A1 and A2 based on the duration of target input, following hierarchical structure of ACtx.

We next computed NCs between conditions for each subregion to investigate trial-by-trial fluctuation as an index of synchronized behavior to encode re-occurring sound. For A1 L4, we only observed an interaction effect between segment duration (short vs. long segment) and condition (Target vs. Random) on NCs (*F*(1,4898) = 10.89, *p* = 0.001). From post-hoc paired t-tests, reduced NCs were observed for Target_rep_ compared to Random_rep_ in A1 L4 (*t*(4898) = −3.06, *p* = 0.012 for Target_rep_ and Random_rep_ pair, *t*(4898) = 1.59, *p* = 0.11 for Target_long_ and Random_long_ pair). In subregions from superficial layers, we observed significant differences for NCs from condition and segment duration, as well as interaction between two factors (A1 L2/3: *F*(1,4651) = 79.42, *p* < 0.0001 for condition, *F*(1,4651) = 27.91, *p* < 0.0001 for segment duration, *F*(1,4651) = 59.92, p < 0.0001 for interaction; A2 L2/3: *F*(1,881) = 25.58, *p* < 0.0001 for condition, *F*(1,881) = 65.65, *p* < 0.0001 for segment duration, *F*(1,881) = 56.90, *p* < 0.0001 for interaction).

We further observed greater NCs for Target_rep_ compared to Random_rep_ only from superficial layers of A1 and A2 (A1 L2/3: *t*(4651) = 9.02, *p* < 0.0001 for Target_rep_ and Random_rep_ pair, *t*(4651) = - 1.85, *p* = 0.12 for Target_long_ and Random_long_ pair; A3 L2/3: *t*(881) = 10.39, *p* < 0.0001 for Target_rep_ and Random_rep_ pair, *t*(881) = 0.5, *p* = 0.34 for Target_long_ and Random_long_ pair). Since NCs are thought to reflect functional connectivity between neurons, these data show that Target stimuli activate highly connected cell pairs that might reflect neuronal ensembles selective for the Target sound, only when the frozen segment is short (180 ms), and only in superficial layers. Together, this suggests functional connectivity starts to emerge from within the superficial layers of ACtx where more active corticocortical projection takes place, compared to thalamocortical recipient layers (L4) of A1.

## Discussion

We studied the early phase of implicit learning of the spectro-temporal stimulus statistics by investigating how the random re-occurrences of complex sounds (DRCs) changes neuronal activity to the repeated DRC. We studied three different ACtx processing areas along the cortical hierarchy: A1 L4, A1 L2/3, and A2 L2/3, using in vivo 2-photon imaging. We observed similar neural dynamics across ACtx areas during implicit learning suggesting that implicit learning is present across ACtx areas even from the earliest ACtx processing stages. While all subregions in ACtx show adaptation of neuronal responses to re-occurring sounds, local networks within subregions differ along the hierarchical structure.

Auditory memory enables us to segregate and understand information from complex auditory streams by identifying relevant features retrieved from prior knowledge. Thus, rapid encoding of newly given sound input by neurons in ACtx, which can be referred to as a ‘learning phase’ of new sound, plays a crucial role to maintain efficient central auditory processing (Rauschecker and Tian, 2000; Zatorre and Belin, 2001; Poremba and Mishkin, 2007; Bendor et al., 2012; Moerel et al., 2012; Atiani et al., 2014). A series of previous studies have established rapid plasticity of neurons in ACtx, especially from adaptation of neuronal responses to a repeatedly presented stimulus in various directions (Rees et al., 1997; Fritz et al., 2003; Nelken, 2014; Parras et al., 2017; Yarden and Nelken, 2017). However, the natural environment is much more complex and unpredictable, thus how encoding processes complex sounds in more naturalistic listening setup remains elusive.

We here show that a subset of neurons in ACtx of mice show neural adaptation to complex stimuli in a passive, implicit learning setting. We adapted an experimental paradigm from human studies that efficiently triggered implicit learning of abstract and complex sounds (e.g., (Agus et al., 2010; Andrillon et al., 2015; Kang et al., 2021). Our findings provide advances in understanding how efficient encoding of sound information is processed in ACtx. In general, a repetitive presentation of a specific stimulus with relatively short inter-stimulus-interval seemed to require observing changes in cell activity to the stimulus. However, response changes of cell activity to re-occurring sounds in ACtx rapidly occurs even within its random, intermittent presentations, as early as its second presentation, with much higher tolerance to any additional “noise” (Random sounds in this case) than we have known (Chew et al., 1995; Chew et al., 1996; Ulanovsky et al., 2004; Nelken, 2014; Yarden and Nelken, 2017; Lu et al., 2018; Yaron et al., 2020).

Our findings show that changes in neuronal responses for encoding a given sound are initiated rapidly, as early as from its second presentation, and is resilient to the timing of its re-occurrences. These findings closely match the ability of human auditory processing (Agus et al., 2010; Snyder and Gregg, 2011; Andrillon et al., 2015; Bianco et al., 2020). Our observation of clear signal difference between Target and Random sounds in even more complex listening settings have been achieved by simultaneous imaging of multiple cells within a relatively large field of view (512 x 512 um). We found that the subset of cells within ACtx that process the encoding of complex sounds seemed to be distinctive from cells that are just responsive to pure tones. 2-photon imaging allowed us to acquire the activity of those cells specifically responsive for complex auditory patterns within the same layer and subregion that are difficult to capture with electrophysiology, while still focusing on each subregion (A1 or A2).

What drives decreased amplitude of excitatory cells for Target conditions still remains an open question. Reduced responsiveness could be due to decreased excitation or increased inhibition. We speculate that inhibitory interneurons likely drive the effect as they play a crucial role in shaping cortical activity (Wehr and Zador, 2003; Kaur et al., 2004; Wu et al., 2008; Isaacson and Scanziani, 2011; Chen et al., 2015; Natan et al., 2015; Natan et al., 2017; Yarden et al., 2022). Particularly, parvalbumin (PV) interneurons in addition to somatostatin (SST) neurons are a candidate for sensory adaptation or actively modulating cortical responses for sensory inputs in different conditions (Kaplan et al., 2016; Maor et al., 2019; Cooke et al., 2020; Yarden et al., 2022). Moreover, close inspection of the DRC traces shows a rise during the last segment, which is post-sequence random DRC segment, as the most noticeable feature from traces in A2 indicating more sustained activity during sound presentation consistent with an effect of inhibition. The rise during post-sequence duration could also be related to offset-related responses (i.e., offset of ‘target’ duration) which have been reported as learning-related activity, especially in high-order regions (Chong et al., 2020; Lee and Rothschild, 2021; Anandakumar and Liu, 2022) to be learning-related activity.

ACtx contains multiple subregions and the hierarchical structure of ACtx has been often reported especially for high-order processing (Lim et al., 2016; Lu et al., 2018). While adaptation of neuronal responses for Target sounds is observed across all subregions in ACtx, differences across subregions also seem to exist. We find that cell types that show distinctive changes in neuronal responses for Target sounds differ across subregions. Although a large proportion of DRC responsive cells (~80% cells) were not responsive to pure tones, we identified subgroups of cells that were responsive to DRC and pure tones. We were able to identify their classic tonal tuning properties and relate tuning properties to learning-related response changes. This analysis revealed that DRC and tonally responsive cells showed different adaptation characteristics across subregions. While in A1 L4, cells with more classic tuning property (e.g., H, I, V-shaped cells) showed similar learning-related response changes, in A1 L2/3 cells with sparse tuning property (specific tuning to a single frequency and sound level) showed learning-related response changes. We speculate that such a finding is due to different local circuits across layers and subregions, potentially from different projection pathways and number of inputs (Oviedo et al., 2010; Hackett, 2011; Meng et al., 2017; Bowen et al., 2020). We consider cells with the classic tuning property are more broadly encoding frequency related information compared to S-type sparse cells (Liu et al., 2019). Then in A1 L4, where thalamocortical projection especially from MGv is the most predominant, cells that process a wider range of spectral information (i.e., cells with classic V-shaped tuning properties) also encode any information that is required for further high-order auditory processing. On the other hand, from A1 L2/3, where corticocortical projections are more predominant, cells process more distinct information from sensory input, to either process tonal information of short sound input or high-order auditory information. Thus, sparse FRA responsive cells in fact mainly play a role in encoding complex sounds rather than shaping tonotopy within the layer and A1. In A2, potentially receiving inputs from both MGv and MGd (Ohga et al., 2018; Liu et al., 2019) along with corticocortical projection, even tonally responsive cells, regardless of its tuning property, are actively processing complex sounds. These considerations suggest that cells in subregions within ACtx may be distinctively characterized depending on information projection circuits (Sołyga and Barkat, 2019; Zempeltzi et al., 2020).

To study this speculation, we further studied the functional connectivity across cells by measuring the population correlations for each subregion. The higher signal correlation (SCs) as an index of response synchrony across cells for Target sounds, regardless of segment duration, were more clearly observed from A2 than in the different layers of A1. Specifically, SCs between Target_rep_ and Random_rep_ conditions in A1 revealed no distinctive neuronal synchrony established for Target sounds. This could be because the short segment duration for Target_rep_, compared to Target_long_, might not be sufficient to establish signal correlation across cells in A1. In fact, information from neural signals can have a better decoding ability when sound contains more information (O’Sullivan et al., 2015). Additionally, no difference of SCs between Target_rep_ and Random_rep_ conditions could be due to repeating patterns embedded within sequence for both Target_rep_ and Random_rep_ thus cells in A1 already shape signal similarity towards the end of the sound in both cases. Increased synchrony that emerged even for Target_rep_ in A1 when the correlation was computed on the trace of only the first segment suggests that it is likely that local networks for re-occurring snippets are established in much shorter timescales in A1. On the other hand, A2 cells have less variability for encoding sound inputs across sound duration, at least between 180 ms and 900 ms, and process sound input as a whole, instead of more fine-tuned spectrotemporal features of shorter segment embedded within the sequence. This finding is parallel to more local temporal processing in A1 and global processing in A2 (Asokan et al., 2021; Kang et al., 2021). Furthermore, greater noise correlation representing functional connectivity was observed only for Target sounds with short segment duration, and only in superficial layers of A1 and A2 and, where more corticocortical projections are predominant. Overall, this indicates that learning of newly present complex sounds is established from a distinctive group of cells with greater connectivity that seem to emerge along the hierarchical structure of ACtx. We speculate that these neurons form an interconnected network, not only within, but also across regions.

Overall findings thus suggest that rapid learning of complex sounds in ACtx is generated by only a fraction of population cells in ACtx with distinctive and shared characteristics across subregions. Its rapid and robust information encoding is in parallel with findings in human auditory perception regarding developing rapid and efficient auditory memory in complex auditory scenes.

## Acknowledgements

HK and POK conceived study. HK performed all experiments and analyzed data. POK and HK wrote the manuscript. Supported by NIH U19NS107464 and NIH R01DC17785.

